# Evolutionary dynamics of the pan-centromere architecture in oaks

**DOI:** 10.64898/2026.11.05.764454

**Authors:** Qiong-Qiong Lin, Yi-Ye Liang, Wei-Shu Yan, Yong-Hang Run, Han-Li You, Biao-Feng Zhou, Shao-Jun Ling, Zhao Shen, Jing-Shu Wang, Jing-Wei Niu, Wen-Ji Luo, Liang-Jing Qiao, Sheng-Chen Shan, Lina Elshaer, Audrey Tucker, Chen Feng, Yong Shi, Rebekah Mohn, Yu Liang, Nai-Li Zhang, Xiao-Juan Liu, Andrew Hipp, Paul S. Manos, Jeannine Cavender-Bares, Pär K. Ingvarsson, Ian Henderson, Douglas E. Soltis, Pamela S. Soltis, Ke-Ping Ma, Zhu-Kuan Chen, Wen-Li Zhang, Baosheng Wang

**Author notes:** These authors contributed equally (Qiong-Qiong Lin, Yi-Ye Liang).

## Abstract

Centromeres ensure faithful chromosome segregation and genome stability, yet their DNA evolves rapidly, presenting a long-standing centromere paradox. Disentangling the evolutionary forces shaping this paradox is crucial for unravelling genome evolution and speciation. Here we characterise the genomic and epigenomic architecture of >1,300 candidate centromeres from 111 high-quality *Quercus* (oak) genomes. We show that centromere turnover is driven by lineage-specific expansion of *QuCEN146* satellite repeats, which displace ancestral LTR-rich sequences into flanking regions. Massive *QuCEN14*6 amplification arose in the ancestor of Eurasian white oaks that experienced continental range expansion, where genetic drift may have promoted fixation of satellite-rich centromeres. Population-scale analyses in *Q. serrata* reveal extensive structural and sequence polymorphism in satellite-rich centromeres, generated by array expansion and contraction, strand-orientation shifts, and interspecific introgression. By contrast, LTR-rich centromeres of *Q. variabilis* exhibit extremely low polymorphism and high collinearity across populations, a pattern sustained by suppressed local recombination, linked selection, and constrained LTR proliferation and deletion. We further demonstrate chromosome-independent evolutionary patterns for both satellite-rich and LTR-rich centromeres in these two species. Collectively, these findings demonstrate how evolutionary processes drive centromere turnover and shape contrasting centromeric sequence landscapes in oaks, with implications for introgression and reproductive isolation in forest trees.

## Main text

Centromeres are the chromosomal domains where CENH3-containing nucleosomes assemble to direct kinetochore formation, ensuring faithful chromosome segregation and genome stability^1,2^. Despite this deeply conserved function, centromeric DNA evolves rapidly, which defines the long-standing centromere paradox^3^. The centromere drive hypothesis provides the dominant framework to explain rapid centromere evolution, in which centromere variants with biased segregation in asymmetric female meiosis acquire a transmission advantage^4–6^. Nonetheless, demographic processes and genetic drift may also contribute to the fixation of novel centromere variants^7^, and their relative contributions remain poorly resolved across species. Because meiotic crossover recombination is suppressed in centromere-proximal regions, divergence in centromeric DNA may accumulate between lineages without being homogenised by gene flow^8,9^. Such divergence may directly compromise chromosome segregation in hybrids and contribute to postzygotic barriers, promoting reproductive isolation and speciation^4,10^. Understanding how centromeres evolve is therefore critical to linking sequence evolution to reproductive isolation, introgression, and speciation^11^.

Plant centromeres are often composed of satellite repeat arrays, long terminal repeat (LTR) retrotransposons, or both^11–20^. Centromere evolution is consequently driven by the dynamics of satellite arrays and LTRs, with the underlying processes and mechanisms differing among species^1,21–31^. For example, satellite homogenisation processes produce higher-order repeats and may efficiently purge invading LTRs in *Arabidopsis*^32^. In contrast, LTR invasion may lead to disruption of ancestral centromeric satellite arrays and facilitate centromere repositioning in rice^33^. These contrasting models raise the question of how the balance between satellite homogenisation and LTR invasion shapes centromere turnover across plant species. Population genomic studies have revealed rapid post-speciation differentiation and high intraspecific variation at centromeres in plants and animals^11,32–36^. However, in domesticated maize and pepper^37,38^, centromeric diversity is markedly reduced on specific chromosomes, a condition that is associated with domestication bottlenecks, selection on centromere-linked genes, and locally suppressed meiotic recombination. Whether reduced centromeric diversity can arise in natural populations, and whether its occurrence is related to centromeric sequence architecture or to local recombination and selection regimes, remain largely unexplored. Addressing these questions requires quantifying centromere variation across both interspecific divergence and intraspecific polymorphism, ideally in a system harbouring both satellite-rich and LTR-rich centromeres.

The genus *Quercus* (oaks) comprises ∼450 species and dominates Northern Hemisphere forests^39–41^, representing an evolutionarily successful woody plant lineage^42^. Oak genomes are highly conserved, with a stable karyotype (*n* = 12) and extensive collinearity even among distantly related species^43–46^. Consistent with this conserved genomic architecture, widespread interspecific introgression has been repeatedly documented among oak species^47–53^. Large satellite arrays have been reported in predicted centromeric regions of sections *Quercus* and *Virentes*^43,54,55^, but not in several other sections examined so far^45,56–58^, suggesting that centromeric sequence composition may differ among oak lineages. Such centromeric variation may reshape recombination landscapes and influence genome-wide patterns of introgression in hybridising lineages^4^. However, the full scope of this variation across oak species remains largely unexplored, because functional centromeres in oaks have not been defined owing to the lack of CENH3 chromatin immunoprecipitation followed by sequencing (ChIP-seq) data. Therefore, it remains unclear how centromeres evolve over millions of years within otherwise highly conserved oak genomes, what evolutionary forces drive strikingly divergent centromere architectures across the genus, and whether such centromeric variation is related to patterns and processes of interspecific hybridization.

Here, we performed comparative and population genomic analyses of 111 high-quality *Quercus* genomes. By combining a sequence-analysis workflow with CENH3 ChIP-seq, DNA methylation, and Hi-C data, we profiled the genomic and epigenomic architecture across more than 1,300 candidate centromeres. We dissected the evolutionary processes and mechanisms driving interspecific centromere turnover and those shaping intraspecific centromeric variation in two representative species with contrasting centromere architectures. Our study provides new insights into the evolutionary dynamics of centromeres within and across oak species.

## Results

### Prediction and validation of Quercus centromeres

To investigate the evolutionary dynamics of centromeres in *Quercus*, we analysed the genomes of 32 oak species, representing all eight recognised sections of the genus (Fig. 1a and Supplementary Table 1). All genomes were assembled using long-read sequencing data, yielding high-fidelity and contiguous assemblies (Supplementary Table 1). To predict candidate centromeric regions, we implemented a CEN-score framework based on hallmark features of plant centromeres, including depleted gene density and enrichment of satellite repeats and/or LTR retrotransposons^59^ (Methods). For each species, predicted centromeric regions were anchored to all 12 chromosomes and displayed high contiguity (Supplementary Fig. 1 and Supplementary Table 1). Of the 384 predicted centromeric regions, 248 (64.6%) were gapless (Supplementary Table 2), and very few showed sequence collapse during genome assembly (Supplementary Fig. 1). A 146-bp satellite monomer, previously reported as a putative centromeric satellite in *Quercus*^43,55^, was consistently detected in all oak species (Supplementary Table 2). We named this ubiquitous satellite repeat *QuCEN146*, which serves as the core target for tracking the evolutionary dynamics of *Quercus* centromeres. For ten species with both methylation and Hi-C sequencing data, candidate centromeres displayed significantly elevated methylation levels and reduced chromatin interaction frequencies relative to chromosome arms (Supplementary Figs. 2-4), consistent with centromeric chromatin properties reported in other species^60–63^. The robustness of the CEN-score approach was further confirmed by an independent algorithm CentIRE^64^ (Supplementary Data 1). Together, we established a high-confidence set of oak centromere candidates, enabling systematic investigation of their composition and evolutionary dynamics across the genus.

**Fig 1:**
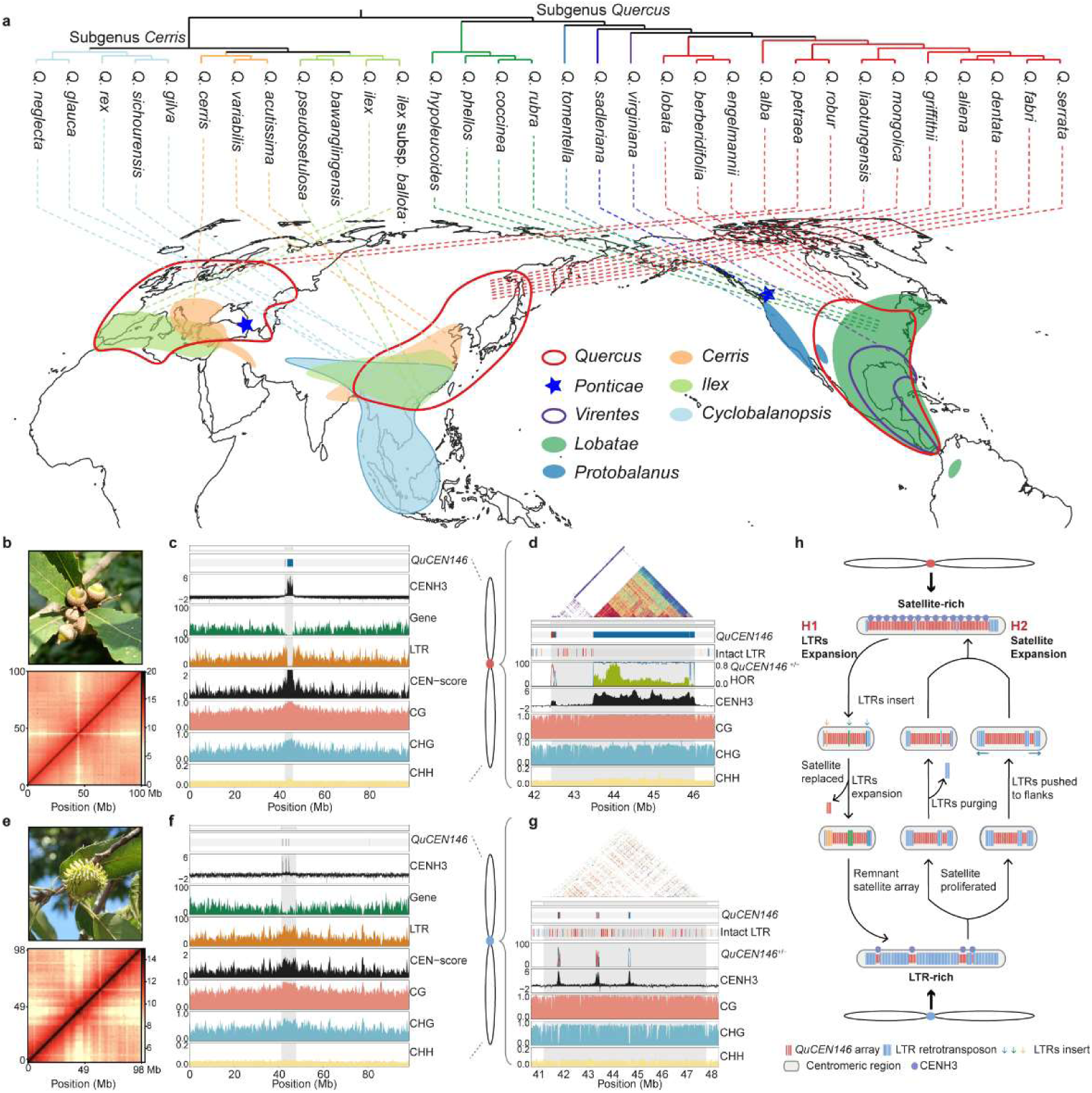
Centromere prediction and characterization in the genus *Quercus*. **A**, Phylogenetic relationships and geographic distributions of 32 oak species representing eight recognized sections of *Quercus*. Top: phylogenetic tree constructed using 1,254 nuclear genes, with branches coloured by section; bottom: global distribution map of the eight *Quercus* sections. **B**, Hi-C chromatin interaction heatmaps for chromosome 2 (bottom) of *Quercus serrata* (top). **C**, Chromosome 2 profiles for *Q. serrata*. From top to bottom: red and blue bars denote *QuCEN146* arrays on the forward and reverse strands, respectively; densities of genes (green) and LTR retrotransposons (yellow); CEN-scores; CENH3 ChIP–seq enrichment (log₂(ChIP/input), black); and DNA methylation profiles in CG (red), CHG (light blue) and CHH (yellow) contexts. Grey shading indicates predicted centromeric regions. All metrics were calculated in 200-kb non-overlapping windows. **D**, Magnified views of centromeric region on chromosome 2 in *Q. serrata*. From top to bottom: StainedGlass sequence identity heatmaps; red and blue bars denoting *QuCEN146* arrays on forward and reverse strands; coloured bars representing different intact LTR families; densities of *QuCEN146* repeats on the forward (red) and reverse (blue) strands; HOR-scores (green); CENH3 ChIP–seq enrichment (log₂(ChIP/input), black); and DNA methylation profiles in CG (red), CHG (light blue) and CHH (yellow) contexts. All metrics were calculated in 10-kb non-overlapping windows. **E**, Hi-C chromatin interaction heatmaps for chromosome 2 of *Quercus variabilis*. **F**, Chromosome 2 profiles for *Q. variabilis*. Layout and all genomic tracks match panel c. **g**, Magnified views of centromeric region on chromosome 2 in *Q. variabilis*. Layout and all genomic tracks match panel d. **h**, Two hypotheses for centromere turnover in oaks. Under hypothesis H1 (LTRs expansion), LTR retrotransposons insert into a satellite-rich centromere and proliferate, progressively replacing *QuCEN146* arrays, leaving only residual satellite arrays. Under hypothesis H2 (satellite expansion), *QuCEN146* arrays proliferate within an LTR-rich centromere, purging LTR retrotransposons or displacing them towards centromere flanks. Red bars represent *QuCEN146* arrays; blue bars represent LTR retrotransposons; coloured arrows denote LTR insertions; grey rounded rectangles represent centromeric regions; light purple dots denote CENH3.

To functionally validate candidate centromeres, we performed CENH3 ChIP–seq on representative species of the two *Quercus* subgenera differing in centromeric repeat content. In *Quercus serrata* (subgenus *Quercus*; Fig. 1b), where predicted centromeres were dominated by *QuCEN146* satellite arrays, CENH3 exhibited continuous enrichment fully overlapping the predicted centromeric domains (Fig. 1c,d and Supplementary Fig. 5). In contrast, in *Quercus variabilis* (subgenus *Cerris*; Fig. 1e), whose candidate centromeres were LTR-rich with limited *QuCEN146* abundance, CENH3 formed discrete punctate peaks (2–4 peaks per chromosome) within each predicted centromeric region (Fig. 1f,g and Supplementary Fig. 6). These CENH3 peaks largely co-localised with sparse *QuCEN146* arrays, whereas the surrounding LTR-rich portions lacked obvious CENH3 enrichment (Fig. 1g and Supplementary Fig. 6). This conserved targeting of CENH3 to *QuCEN146* arrays corresponded well with the overall conservation of oak CENH3 protein (Supplementary Fig. 7). Notably, CENH3-enriched domains in both species were depleted of CHG methylation (Fig. 1d,g and Supplementary Figs. 5-6), a pattern also observed in *Arabidopsis*^32,63^. Collectively, our ChIP–seq data demonstrated that CENH3 preferentially associated with *QuCEN146* satellite arrays, yet CENH3 binding profiles differed markedly between *Q. serrata* and *Q. variabilis*, corresponding to their contrasting centromeric repeat landscapes. These findings identify functional centromeric regions in oak species and further support the reliability of our CEN-score framework for genome-wide centromere prediction.

### Satellite expansion remodels centromeric landscapes in Quercus

Despite extensive collinearity across oak genomes, predicted centromeric regions exhibit considerable interspecific heterogeneity in sequence composition (Supplementary Table 3 and Supplementary Fig. 8). In subgenus *Quercus*, candidate centromeres of sections *Quercus* and *Ponticae* are predominantly composed of the satellite repeat *QuCEN146*, which constitutes 80–90% of centromeric sequences (Fig. 2a,b and Supplementary Table 4). The only exceptions are three North American white oak species with low-to-moderate *QuCEN146* copy numbers (Supplementary Table 2). In contrast, centromeric regions in sections *Cerris* and *Cyclobalanopsis* of subgenus *Cerris* are dominated by LTRs (50-70% of centromeric sequences) with very low *QuCEN146* abundance (<3%; Fig. 2a,b and Supplementary Table 4). The remaining oak species possess centromeres dominated by LTRs but with moderate *QuCEN146* abundance (Fig. 2a,b and Supplementary Table 4). This heterogeneous sequence composition reflects centromere turnover: transitions between LTRs and *QuCEN146* repeats within oak centromeres.

**Fig 2:**
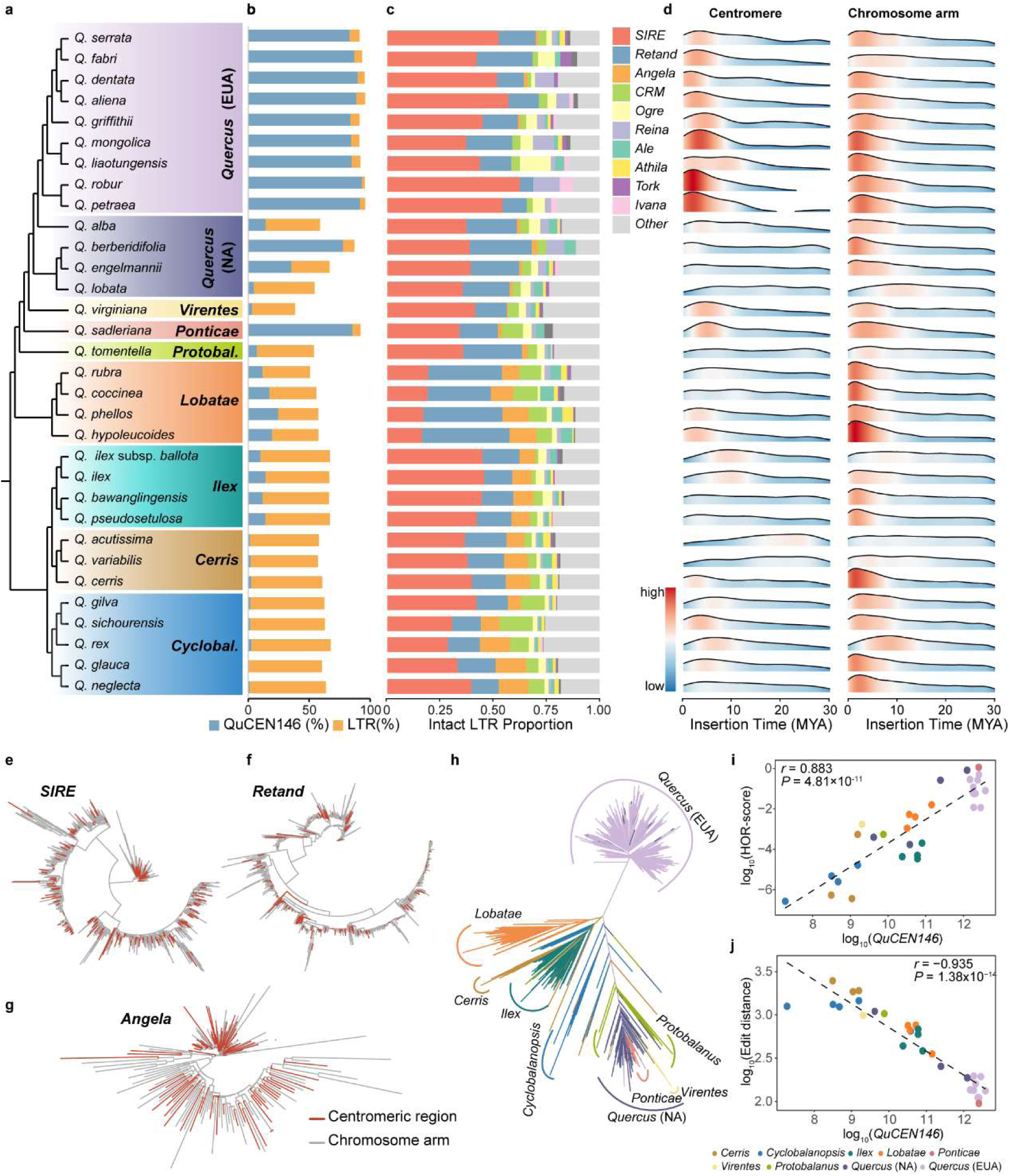
Divergent centromeric composition and *QuCEN146* satellite dynamics across oaks. **a**, Phylogenetic relationships of 32 *Quercus* species spanning eight sections. Eurasian (EUA) and North American (NA) lineages of section *Quercus* were coloured differently. **b**, Bar charts showing *QuCEN146* and LTR retrotransposon density in candidate centromeric regions across species. **c**, Relative proportions of the top 10 most frequent LTR retrotransposon families within candidate centromeric regions; remaining families are grouped as “Other”. **d**, Heatmaps showing insertion time distributions of intact LTR retrotransposons from centromeres (left) and chromosome arms (right). Darker regions indicate higher density. MYA, millions of years ago. **e–g**, Phylogenetic trees of the three most abundant centromeric LTR families (*SIRE*, *Retand*, *Angela*) from *Q. variabilis*, whose candidate centromeric regions are LTR-rich. Centromeric LTR sequences (red) are interspersed with those from chromosome arms. Similar patterns were observed in other oak species with LTR-rich centromeres (Supplementary Fig. 11). **h**, Phylogenetic relationships of non-redundant *QuCEN146* sequences sampled across 32 oak species. Branches are coloured by section, with EUA (light purple) and NA (dark purple) geographic lineages highlighted within section *Quercus*. **i,j**, Correlations between HOR score and *QuCEN146* abundance (k), and between edit distance and *QuCEN146* abundance (l). All variables are log_10_-transformed. Each dot represents one species, and is coloured by section and geographic lineage.

We propose two plausible hypotheses to explain the diversity of *Quercus* centromere: either LTRs expanded and replaced ancestral satellite repeats (H1), or satellite repeats proliferated, purging pre-existing LTR retrotransposons, or displacing them to flanking regions (H2; Fig. 1h). To distinguish between these hypotheses, we systematically profiled centromeric LTRs in *Quercus*. Multiple lines of evidence argue against recent LTR bursts in oak centromeres. First, LTR-rich centromeres exhibited weak signals of recent LTR insertion, with the majority of intact centromeric LTRs having insertion ages >5 million years ago (MYA; Fig. 2d and Supplementary Fig. 9). Second, the ratio of intact-to-solo LTRs in centromeric regions was comparable to that on chromosome arms (Supplementary Table 4), supporting similar long-term evolutionary dynamics rather than recent centromere-specific proliferation. Third, LTR-family compositions were similar between centromeres and chromosome arms (Fig. 2c and Supplementary Fig. 10), and centromeric LTRs were interspersed with arm-located LTRs across the phylogenetic tree for the two most abundant LTR families (*SIRE* and *Retand*) (Fig. 2e,f and Supplementary Fig. 10). Notably, *Angela* retrotransposons were specifically enriched in centromeres relative to chromosome arms (∼10% vs. <1%) in subgenus *Cerris* and section *Lobatea* (subgenus *Quercus*) (Fig. 2c and Supplementary Fig. 11). Nonetheless, centromeric *Angela* elements lacked signals of recent bursts (Fig. 2g and Supplementary Figs. 9,11), representing a subtle centromere-preferential accumulation rather than recent centromere-specific expansion. Collectively, these findings rule out recent LTR bursts in centromeres, suggesting that *QuCEN146* repeat expansion may have driven centromere turnover in oaks.

We further dissected the expansion dynamics of *QuCEN146* across oak lineages. Phylogenetic analysis of non-redundant *QuCEN146* sequences resolved two major clades: the Eurasian white oak (EUA) clade and the non-EUA clade encompassing all other oaks, including North American white oaks (NA) (Fig. 2h). Comparison of consensus sequences revealed substantial divergence between the *QuCEN146* repeats of these two clades (Supplementary Fig. 12). These results suggest at least two major amplification events of *QuCEN146*, corresponding to the EUA and NA lineages. Within the EUA clade, *QuCEN146* sequences from different species intermingled on the phylogenetic tree, consistent with a shared ancestral expansion event (Supplementary Figs. 12-13). In contrast, NA sequences tended to cluster by species, suggesting independent expansion followed by subsequent divergence. Beyond these two lineages, *Quercus sadleriana* (section *Ponticae*) carries an exceptionally high *QuCEN146* copy number with high sequence similarity to NA (Fig. 2h and Supplementary Figs. 12-13), consistent with their ancient hybridization^65^. In the remaining non-EUA lineages, *QuCEN146* sequences tended to cluster by section, consistent with long-term satellite stability and sequence divergence (Fig. 2h, Supplementary Fig. 13). Collectively, we indicate large-scale *QuCEN146* amplification primarily in the EUA lineage, alongside more isolated cases in certain NA species and a distinct instance in *Q. sadleriana*.

Amplification of centromeric satellites often generates higher-order repeats (HORs)^63,66,67^. Therefore, we quantified HOR-scores (HOR number per basic repeat unit)^32^ and edit distance in predicted centromeres to assess the activity of *QuCEN146* arrays. We found that *QuCEN146* copy number was positively correlated with HOR-scores (Spearman’s p = 0.88, *P* = 4.81×10^-11^) and negatively correlated with edit distances (Spearman’s p = -0.94, *P* = 1.38×10^-14^; Fig. 2i,j). For example, the satellite-rich centromeres of the EUA lineage had HOR-scores (0.144-0.743) approximately four-fold higher than the LTR-rich centromeres of subgenus *Cerris* (0–0.038; Fig. 2i, Supplementary Fig. 14 and Supplementary Table 2). This concordance between *QuCEN146* expansion activity and satellite-rich centromere architecture, together with the absence of recent centromeric LTR expansion, supports satellite proliferation as the primary driver of centromeric turnover in oaks. Furthermore, flanking regions of the centromeric array in *Q. serrata* showed high collinearity with the LTR-rich portion of centromeric regions in *Q. variabilis* (Supplementary Fig. 15). This result suggests that satellite expansion displaced ancestral LTR-rich centromeric sequences towards satellite flanks rather than eliminating the ancestral centromeric compartment.

### Extensive diversification of satellite-rich centromeres in Quercus serrata

To characterise the patterns and underlying mechanisms driving centromeric variation, we focused on *Quercus serrata*, a Eurasian white oak with satellite-rich centromeres. We generated 46 high-quality genomes for this species (Fig. 3a and Supplementary Table 5) and predicted 552 high-confidence centromeric regions (Supplementary Fig. 16, Supplementary Table 6 and Supplementary Data 2). Across centromeric regions, we identified 10,922,054 copies of *QuCEN146* repeats, 27.6% of which were singletons (Supplementary Table 7). The number of non-redundant *QuCEN146* sequences continued to rise with increased sampling (Supplementary Fig. 17a), revealing substantial monomer-level variation within this satellite family. All candidate centromeres of *Q. serrata* were dominated by long *QuCEN146* satellite arrays with extensive sequence polymorphism (Supplementary Table 7). We also observed substantial array size variation among chromosomes and individuals, with significantly larger arrays found in western populations (*P* = 3.03× 10^-11^, Mann-Whitney *U* test; Fig. 3b and Supplementary Fig. 17b).

**Fig 3:**
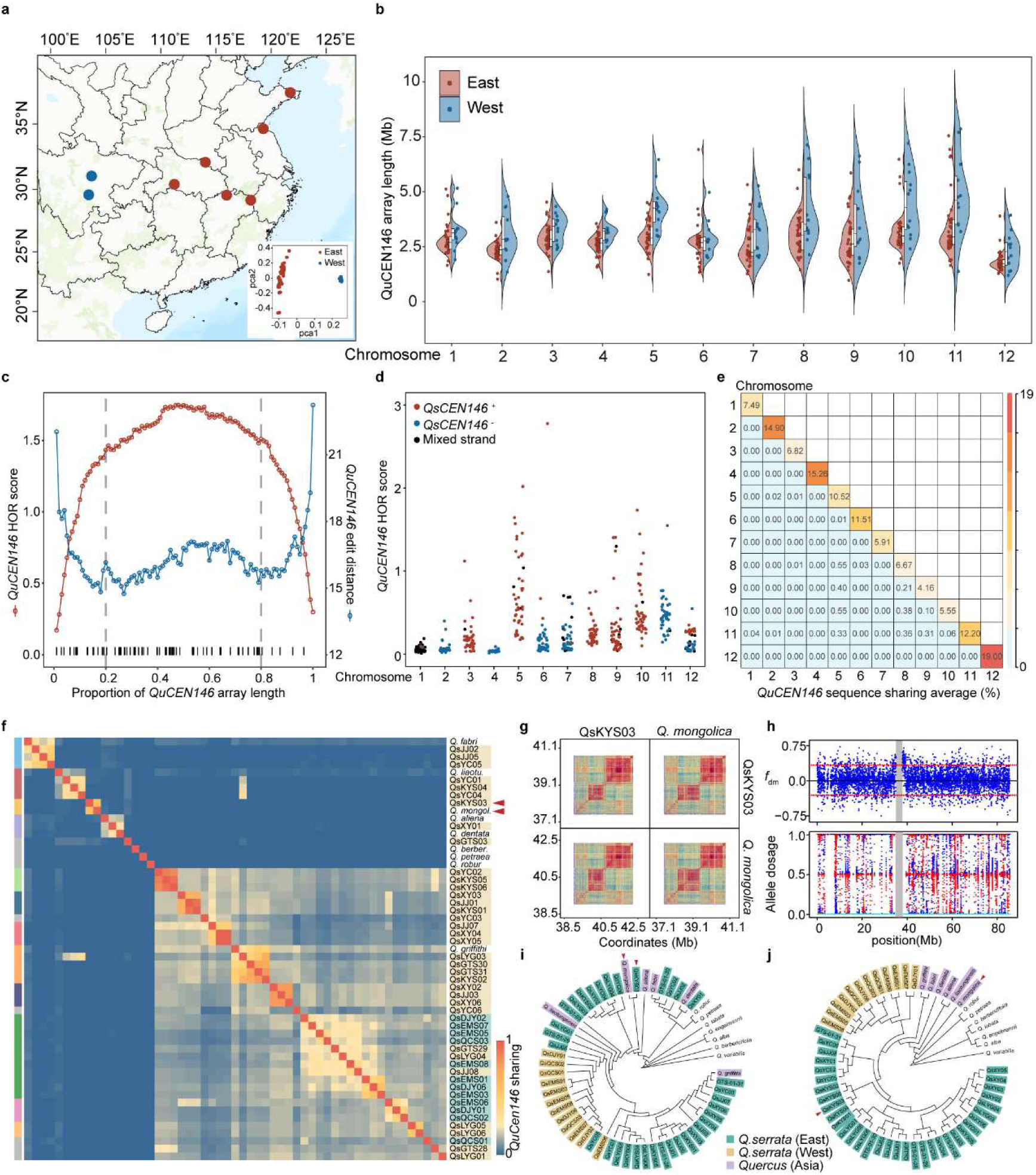
*QuCEN146* satellite variation and centromeric introgression in *Quercus serrata*. **a**, Geographic locations of eight sampled *Q. serrata* populations, comprising 46 accessions (5-6 individuals per population) used for centromere analyses. Populations are coloured according to east (red) and west (blue) genetic groups, defined via principal component analysis (PCA) of genome-wide SNPs. **b**, Length variation of centromeric *QuCEN146* arrays for each chromosome across the 46 *Q. serrata* accessions. Each dot represents an individual, coloured by east (red) or west (blue) genetic group. **c**, Plot of *QuCEN146* HOR scores (red) and edit distances (blue) across normalized length of the satellite arrays. Black ticks denote ultra-long HORs (>40 kb). **d**, HOR scores of centromeric *QuCEN146* arrays per chromosome across 46 *Q. serrata* accessions. Coloured points represent individual centromeres dominated by forward-strand (red) or reverse-strand (blue) arrays (>90% of array). Black points indicate centromeres with mixed strand orientations, where neither strand exceeds the 90% threshold. **e**, Heatmap showing the average proportion of shared identical *QuCEN146* sequences in pairwise chromosome comparisons across the 46 *Q. serrata* accessions. **f**, Heatmap showing the percentage of shared *QuCEN146* sequences between pairwise accessions for chromosome 5. Centromere haplotype (CenHap) groups are indicated on the left, and accession names are labelled on the right. *Q. serrata* accessions are coloured according to east (yellow) and west (green) genetic groups, while other Eurasian white oaks are uncoloured. Red arrows indicate the pair of accessions showing centromeric introgression on chromosome 5. **g**, StainedGlass sequence identity heatmaps illustrating centromeric sequence identity within and between QsKYS03 (*Q. serrata*) and MESQm (*Q. mongolica*). **h**, Signatures of interspecific gene flow between QsKYS03 (*Q. serrata*) and MESQm (*Q. mongolica*) along chromosome 5. Top: *f*_dm_ statistics calculated from ABBA–BABA tests in 10 kb non-overlapping windows; red dashed lines denote the 5th and 95th percentiles of genome-wide *f*_dm_ values. Bottom: allele dosage for QsKYS03 inferred with ELAI, using QsEMS08 (blue), MESQm (red) and SDTS (*Q. variabilis*, light-blue) as source panels. Grey shading indicates predicted centromeric regions. **i,j** Phylogenetic trees of *Q. serrata* and other white oaks, constructed using SNPs in the 500 kb centromere-flanking region of chromosome 5 (i) and all SNPs across this chromosome (j). Trees are rooted with *Q. variabilis*, and branches are coloured by *Q. serrata* genetic groups. Red arrows indicate the pair of accessions showing centromeric introgression on chromosome 5.

To dissect the fine-scale centromere organization, we characterised the HOR structure within *QuCEN146* arrays. Across all candidate centromeres, HORs covered 71.7–99.5% of the total satellite array length (Supplementary Table 8). The core regions of *QuCEN146* arrays exhibited elevated HOR-scores and reduced edit distances relative to the margins, and harbored 60 of the 82 ultra-long HORs (>40 kb; Fig. 3c and Supplementary Fig. 18). This indicates an intense homogenization process within the central regions, consistent with observations in *Arabidopsis*^32^ and rice^33^. Additionally, the degree of sequence homogenization varied substantially among chromosomes (Fig.3d, Supplementary Fig. 19 and Supplementary Table 8), suggesting chromosome-specific homogenization dynamics. In line with such independent evolution of centromeric *QuCEN146* arrays, we observed chromosome-specific mutation spectra and limited inter-chromosomal sharing of these repeats (Fig. 3e and Supplementary Fig. 20).

We further clustered *QuCEN146* arrays into distinct centromere haplotype (CenHap) groups for each chromosome based on pairwise sequence similarity. The CenHap grouping patterns varied drastically among chromosomes and did not correspond to the east-west genetic structure revealed by genome-wide SNPs (Fig. 3f and Supplementary Fig. 21). Despite generally low CenHap sequence similarity among oak species, we detected interspecific introgression that produced highly similar *QuCEN146* array between *Q. serrata* and other Eurasian white oak species. For example, one individual each from *Q. serrata* and *Q. mongolica* fell into the same Chr5 CenHap group (Fig. 3f), reflecting trans-centromeric introgression (Fig. 3g-j). Such centromeric introgression was often restricted to individual chromosomes, rather than affecting multiple chromosomes simultaneously (Supplementary Fig. 21), suggesting that the introgressed centromeric segment has persisted in the *Q. serrata* genome long after ancient hybridization.

We also observed widespread strand-orientation polymorphism of *QuCEN146* arrays across chromosomes and individuals (Fig.4a). This strand-orientation variation is driven by two distinct mechanisms. On Chr12, 22 eastern accessions were assigned to the same CenHap clade and showed elevated HOR-scores and reduced edit distances relative to other samples (Fig. 4b,c). Among them, 18 harboured forward-strand *QuCEN146* arrays, whereas the remaining four carried reverse-strand arrays. Collinearity analysis revealed that these two array types differ by a large-scale trans-centromeric inversion (Fig. 4d and Supplementary Fig. 22), which may have rearranged the orientation of embedded *QuCEN146* arrays. This Chr12 CenHap was absent from all other Eurasian white oaks (Fig. 4b), making it unlikely to have arisen from ancestral polymorphism or interspecific introgression. Therefore, we favour a single-origin scenario for Chr12 CenHap: one ancient inversion produced the rearranged CenHap, which subsequently spread through populations via gene flow (Fig. 4g). In contrast, forward *QuCEN146* arrays on Chr6 exhibit multiple independent origins. Three eastern individuals carried forward-strand arrays but belonged to distinct Chr6 CenHap groups with low sequence similarity (Fig. 4e and Supplementary Fig. 21). These three forward-strand *QuCEN146* arrays exhibited elevated HOR-scores and reduced edit distances relative to reverse-strand arrays (Fig. 4f and Supplementary Fig. 19), consistent with forward-strand expansion driven by extensive tandem duplication and sequence homogenization. These events likely started with local reverse-to-forward strand flipping, followed by rapid duplication and expansion to form large forward-strand arrays (Fig. 4g). Together, our results show that centromeric arrays evolve via distinct mechanisms across chromosomes, reinforcing the chromosome-independent evolution of centromeres.

**Fig 4:**
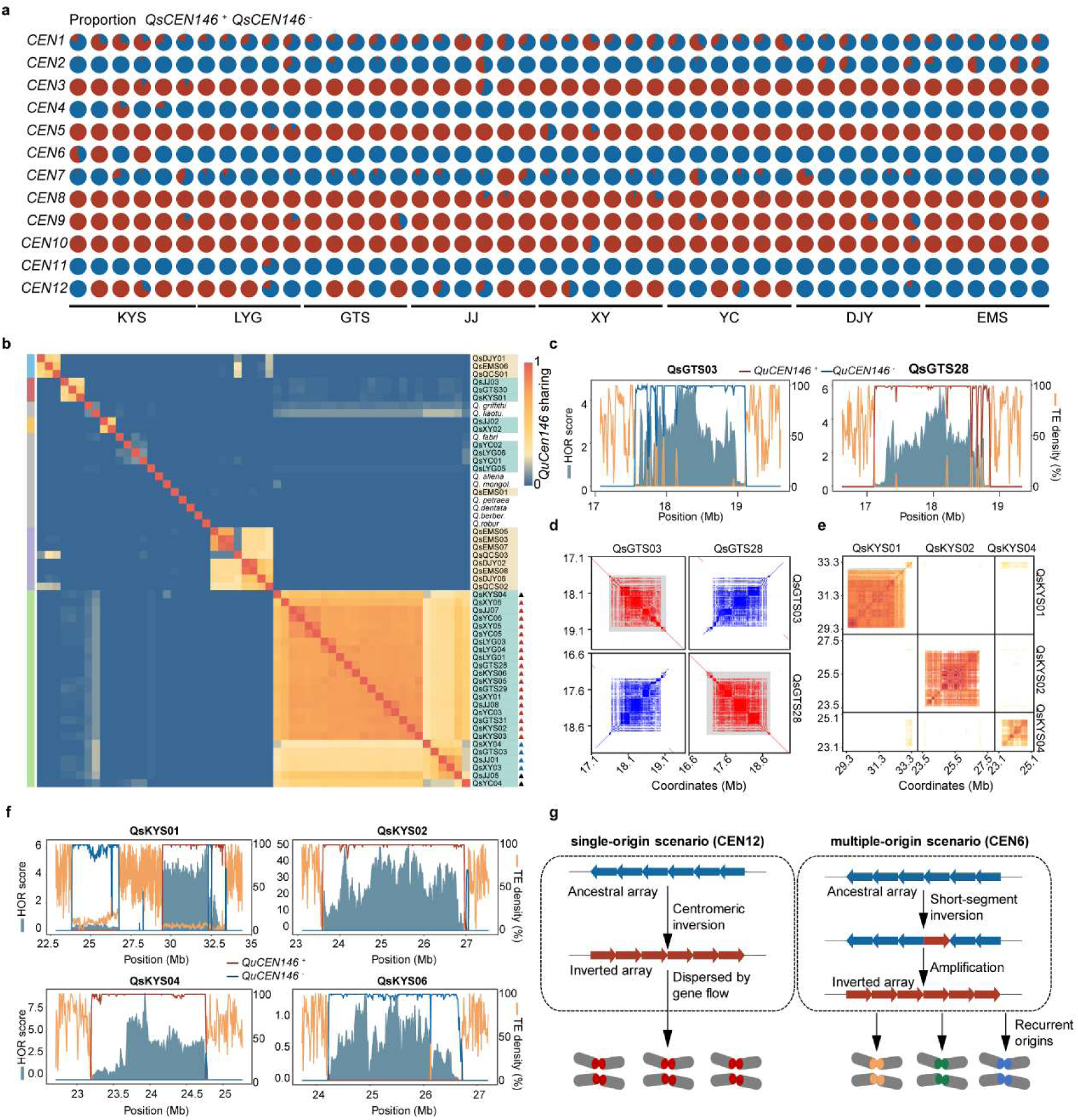
Dynamics of *QuCEN146* satellite array in *Quercus serrata*. **A**, Pie charts showing the proportion of *QuCEN146* array on forward (red, +) or reverse (blue, –) strands for each chromosome (rows) and accession (columns). The eight source populations are labelled at the bottom. **B**, Heatmap showing the percentage of shared *QuCEN146* sequences between pairwise accessions for chromosome 12. Centromere haplotype (CenHap) groups are indicated on the left, and accession names are labelled on the right. *Q. serrata* accessions are coloured according to east (yellow) and west (green) genetic groups, while other Eurasian white oaks are uncoloured. Arrows indicate 18 forward-strand-dominated (red), four reverse-strand-dominated (blue) and three mixed-strand (black) individuals. **C**, HOR scores, *QuCEN146* array density and transposable element (TE) density plotted along chromosome 12 centromeric regions (CEN12) of QsGTS03 (left) and QsGTS28 (right). **D**, Sequence identity dot plot comparing CEN12 in QsGTS03 and QsGTS28 using 100-bp windows, showing a large inversion that reverses the strand orientation of embedded *QuCEN146* arrays. Red and blue indicate similarity between the same and opposite strands, respectively. **E**, StainedGlass sequence identity heatmaps of forward-strand arrays from QsKYS01, QsKYS02 and QsKYS04, showing low similarity consistent with the independent origins of their chromosome 6 CenHaps. **F**, HOR scores, *QuCEN146* array density and TE density plotted along the centromeric regions of four independent chromosome 6 CenHaps: three independently arisen forward-strand haplotypes (QsKYS01, QsKYS02, QsKYS04) and one reverse-strand haplotype (QsKYS06) for comparison. **G,** Schematic models of centromeric array evolution. Single-origin scenario (CEN12): an ancestral inversion produced inverted arrays (blue arrows); this rearranged CenHap subsequently spread across the population via gene flow, as exemplified for chromosome 12 in *Q. serrata*. Multiple-origin scenario (CEN6): independent local inversions and recurrent amplification repeatedly produced inverted arrays (blue arrows) with distinct genetic backgrounds, as exemplified for chromosome 6.

Centromeric satellite arrays are frequently invaded by LTRs^32^. We detected 335 intact and 1,059 soloLTRs within *QuCEN146* arrays, occupying 0.23% of their sequence (Supplementary Table 9). The intact-to-solo LTR ratio was higher inside *QuCEN146* arrays than outside (0.32 vs. 0.18; Supplementary Fig. 23a and Supplementary Table 9), and intact LTRs within arrays were significantly younger than those on chromosome arms (*P* = 2.2× 10^-16^, Mann-Whitney *U* test; Supplementary Fig. 23b). Permutation tests showed that intact LTRs were enriched in array regions with reduced HOR-scores (Supplementary Fig. 23c). These results suggest that LTRs are preferentially purged from *QuCEN146* arrays, a process that likely maintains the structural and functional integrity of *Q. serrata* centromeres. Nevertheless, a subset of LTRs exhibits chromosome-specific accumulation within *QuCEN146* arrays, with only 1-4 copies per array (Supplementary Tables 10-11). This differential LTR purging and retention contributes to centromeric genetic variation.

### Conserved architecture of LTR-rich centromeres in Quercus variabilis

We further extended our population-scale analysis to *Quercus variabilis*, whose centromeres were characterised by a high LTR density and low *QuCEN146* abundance. We predicted 396 high-confidence candidate centromeres from 33 high-quality *Q. variabilis* genomes (Fig. 5a and Supplementary Table 12). In contrast to the highly variable architecture of satellite-rich centromeres in *Q. serrata*, the predicted centromeric regions of *Q. variabilis* displayed remarkable structural and sequence conservation. All candidate centromeres were dominated by LTRs (50–60% of centromeric sequences), with *QuCEN146* satellites restricted to 2-4 small arrays (<3%) (Supplementary Tables 13-14). For each chromosome, the length and sequence composition of its centromeric region, together with the position of *QuCEN146* arrays, were almost consistent across all individuals (Supplementary Table 14 and Supplementary Fig. 24). In line with this compositional stability, centromeric regions exhibited significantly higher collinearity than chromosome arms (SynDiv mean= 0.09 vs. 0.15; *P* = 1.14× 10^-53^, Mann-Whitney *U* test), with even the *QuCEN146* arrays showing near-perfect collinearity (Fig. 5b-e and Supplementary Fig. 25).

**Fig 5:**
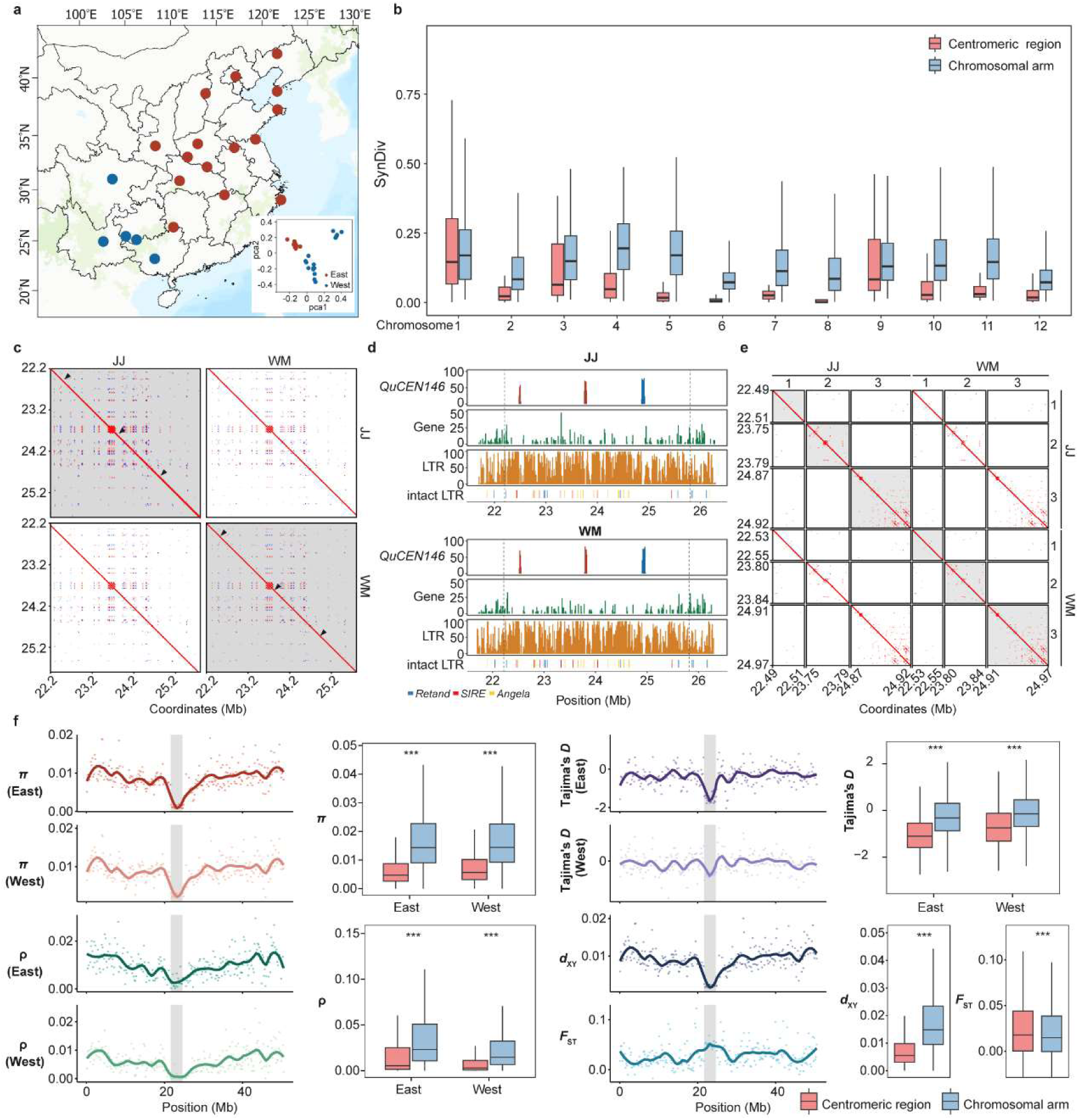
Architecture and genetic variation of LTR-rich centromeres in *Quercus variabilis*. **a**, Geographic locations of 33 *Q. variabilis* individuals from 21 populations across its natural distribution range. Populations are coloured according to east (red) and west (blue) genetic groups, defined by principal component analysis (PCA) of genome-wide SNPs. **b**, Boxplots of synteny diversity (SynDiv) comparing centromeric regions (red) and chromosome arms (blue) across the 12 chromosomes of 33 *Q. variabilis* accessions. Centromeres exhibit significantly higher collinearity (lower SynDiv) than chromosomal arms (*P* < 0.001, Mann–Whitney *U* test). **c–e**, Comparison of centromeric structure on chromosome 8 between two *Q. variabilis* accessions from distinct genetic groups: JJ (east) and WM (west). **c**, Sequence identity dot plots showing high centromeric collinearity between JJ and WM. Black arrows denote the three centromeric *QuCEN146* arrays in each accession. **d**, Densities of *QuCEN146* satellite arrays, genes and LTRs across the centromeric region for JJ (top) and WM (bottom). Black dashed lines indicate predicted centromeric boundaries. **e**, Sequence identity dot plots showing pairwise comparisons of the three *QuCEN146* arrays within centromeric region between JJ and WM. **f**, Comparison of five population-genetic summary statistics between centromeric regions (red) and chromosome arms (blue) on chromosome 6. Each panel shows a Manhattan plot of statistics calculated in 10-kb non-overlapping windows along the chromosome with a loess trend line, paired with a boxplot for group comparisons. Grey shading marks the predicted centromeric region. Significance for each comparison was assessed using the Mann–Whitney *U* test (\*\*\**P* < 0.001).

We next investigated how distinct mechanisms may have contributed to this extreme structural conservation of *Q. variabilis* centromeres. First, the intact-to-solo LTR ratio was lower inside centromeric regions than outside, and intact LTRs within centromeres were significantly older than those on chromosome arms (*P* < 2 × 10^-7^, Mann-Whitney *U* test; Supplementary Table 15 and Supplementary Fig. 26), suggesting low recent LTR proliferation within centromeric regions. Additionally, solo LTRs within centromeres were longer (*P* < 7 × 10^-5^, Mann-Whitney *U* test) and displayed higher CG methylation (*P* = 8.1 × 10^-16^, Mann-Whitney *U* test; multiple linear regression) than their chromosome arm counterparts (Supplementary Tables 15-16), suggesting that localised epigenetic modification may have suppressed the truncation of centromeric solo LTRs. Finally, the *QuCEN146* arrays exhibited extremely low HOR-scores (0–0.02) and high edit distances (20.59–47.50; Supplementary Table 17), indicating long-term satellite inactivity. Altogether, suppressed recent LTR insertions, coupled with constrained solo LTR truncation and quiescent *QuCEN146* satellites, likely maintained the conserved architecture of centromeres in *Q. variabilis*.

Beyond their extreme structural conservation, *Q. variabilis* centromeres also harboured markedly low genetic variation. Nucleotide diversity (π) within centromeric regions was approximately one-third of that on chromosome arms (0.004 vs. 0.012), accompanied by an excess of low-frequency alleles (lower Tajima’s *D*), as well as reduced recombination rate (ρ) and absolute divergence (*d*_XY_) (all comparisons *P* < 1 × 10^-9^, Mann-Whitney *U* test; Fig. 5f, Supplementary Fig. 27 and Supplementary Table 18). These patterns of reduced genetic variation were consistent across all chromosomes and persisted after the removal of centromeric genes and their 1-kb flanking regions (Supplementary Table 18), suggesting that selection on functional genes alone is insufficient to explain the depletion of centromeric diversity. Accordingly, partial correlation analyses within centromeric regions revealed that π was positively correlated with recombination rate (Spearman’s partial ρ = 0.16 - 0.18, *P* < 2.2 × 10^-16^, controlling for gene density; Supplementary Fig. 28). The role of recombination suppression in shaping centromeric diversity was further supported by low nucleotide diversity within gene-poor intervals of *Q. serrata* centromeres, despite high variation in *QuCEN146* arrays (Supplementary Table 18). Collectively, these results suggest that recombination suppression is a major contributor to the reduced genetic variation in *Q. variabilis* centromeres.

Although the centromeres of *Q. variabilis* exhibited high sequence and structural conservation, we found evidence for distinctive evolutionary dynamics across chromosomes. Population structure analyses using centromeric SNPs failed to recover the east–west population divergence pattern delineated by genome-wide SNPs but instead revealed chromosome-specific clustering patterns that varied among chromosomes (Supplementary Figs. 29-30). This observation suggested chromosome-independent evolutionary trajectories for the LTR-rich centromeres of *Q. variabilis*.

## Discussion

The genus *Quercus*, which harbours both LTR-rich and satellite-rich centromeres, provides a powerful system to dissect centromere turnover and its underlying mechanisms. Our study reveals a centromere evolution model driven by satellite expansion: proliferating *QuCEN146* arrays displace ancestral LTR-rich centromeric sequences into flanking regions, where these LTR-rich compartments remain stable over long evolutionary timescales (Fig. 1h). This contrasts with the retrotransposon-induced paradigm described in rice, in which LTR insertions drive centromere turnover and CENH3 nucleosomes assemble on both canonical satellites and centrophilic LTRs^33^. In oaks, however, CENH3 localises exclusively to *QuCEN146* arrays, with no detectable CENH3 enrichment across surrounding LTR-rich regions. Therefore, oak centromeric LTRs appear to constitute an ancient genomic backbone rather than permissive substrates for CENH3 nucleosome assembly.

The rapid evolution of centromeres is widely explained by the centromere drive hypothesis, whereby larger repeat arrays gain a transmission advantage in asymmetric female meiosis^3,68^. However, drive alone cannot fully account for centromere turnover in oaks. First, *QuCEN146* occurs across all *Quercus* sections, indicating that this satellite had already arisen in the common ancestor of oaks by ca. 56 MYA^41^. Yet massive *QuCEN14*6 amplification occurred in only a small subset of lineages, suggesting that centromere drive alone is insufficient to explain the heterogenesous fixation of greatly expanded satellite arrays across oak lineages. Additionally, the most striking *QuCEN146* expansion is restricted to Eurasian white oaks, which experienced a Miocene range expansion out of North America^41,65^. Strong bottlenecks associated with this continental dispersal may have promoted the fixation of satellite-rich centromeres by genetic drift in Eurasian white oaks. Demographic history has also been proposed to contribute to centromeric satellite turnover in *Arabidopsis*^7^.

We further reveal contrasting patterns of genetic variation between satellite-rich and LTR-rich centromeres. In *Q. serrata*, we detected extensive structural and sequence polymorphism in satellite-rich centromeres, arising from array expansion and contraction, strand-orientation changes, and interspecific introgression. This high centromeric variation aligns with observations in other species with satellite-rich centromeres^11,32–34,36,69^ and reflects a phenomenon central to the centromere paradox^3^. In contrast, the LTR-rich centromeres of *Q. variabilis* exhibit extremely low polymorphism and high collinearity across populations. Our results suggest that this centromeric stability is underpinned by regional recombination suppression, linked selection, and reductions in both LTR proliferation and deletion. Reduced centromeric diversity has been reported at specific centromeres of maize and pepper, with this reduction largely attributed to domestication bottlenecks and selection on centromere-linked genes^37,38^. However, despite retaining a large effective population size after diverging from its sister species^56^, *Q. variabilis* showed similarly low diversity across all candidate centromeres. These findings suggest that low centromeric diversity can be sustained in outcrossing populations with high genome-wide diversity, and that this depletion is shaped mainly by a distinct genomic context rather than by direct selection on centromeric genes.

Lastly, we show chromosome-independent evolutionary patterns for both satellite-rich and LTR-rich centromeres in *Quercus*, consistent with observations in other plants^32,33^ and in humans^34^. Because centromeric regions experience low recombination and evolve largely independently of chromosome arms, they may retain distinct signals of hybridization^33,70^ and selection^37,38,71^. By integrating interspecific comparisons and population genomic analyses of centromeres, our study dissects the evolutionary mechanisms driving centromere turnover and shaping contrasting centromeric sequence landscapes in *Quercus*. Future studies combining functional data with complementary analytical approaches will advance our understanding of how centromeric dynamics modulate recombination, introgression and speciation.

## Methods

### Taxon and population sampling

To systematically dissect centromeric variation across a broad phylogenetic continuum of divergence, we sampled 32 *Quercus* species, representing all eight recognized sections of the genus^40^. Our sampling included sections *Quercus* (13 species), *Virentes* (1 species), *Ponticae* (1 species), *Protobalanus* (1 species), and *Lobatae* (4 species) from subgenus *Quercus*; and sections *Ilex* (4 species), *Cerris* (3 species), and *Cyclobalanopsis* (5 species) from subgenus *Cerris*. Species were selected to maximize taxonomic coverage of the major clades resolved in recent phylogenomic studies^41,65,72^. Of these 32 species, 13 were newly sequenced and assembled in the present study, whereas the remaining 19 high-quality genome assemblies were retrieved from publications^45,54–57,73–80^ and the Darwin Tree of Life Project (https://www.darwintreeoflife.org/; Supplementary Table 1). All 19 retrieved assemblies were generated using long-read sequencing data: two used standard PacBio CLR reads, and the remainder used PacBio HiFi or ONT reads; 18 were anchored to chromosomes via Hi-C or Omni-C scaffolding (Supplementary Table 1).

To further characterise centromeric variation at the within-population scale, we focused on two representative species with contrasting centromeric architectures: *Quercus serrata* (subgenus *Quercus*), which harbours satellite-rich centromeres; and *Quercus variabilis* (subgenus *Cerris*), which has LTR-rich centromeres. For *Q. serrata*, we sampled 46 individuals, including 34 from eastern populations and 12 from western populations, capturing the east–west geographic gradient across its distribution range (Supplementary Table 5). For *Q. variabilis*, 17 and 16 individuals were sampled from eastern and western genetic groups, respectively (Supplementary Table 12). Twenty-two *Q. variabilis* assemblies were retrieved from ref^56^, and the remaining 11 individuals were newly assembled for this study (Supplementary Table 12).

### Genomic DNA and RNA extraction, PacBio HiFi, Hi-C and transcriptome sequencing

For all newly sequenced samples, high-molecular-weight genomic DNA was extracted from flash-frozen leaf tissue using the CTAB (cetyltrimethylammonium bromide) method^81^. DNA integrity and contamination were evaluated by 1% agarose gel electrophoresis, and DNA concentration was quantified on a Qubit 2.0 Fluorometer (Life Technologies). PacBio HiFi sequencing libraries were constructed from 15 µg of purified DNA per sample using the SMRTbell Express Template Prep Kit 2.0 (Pacific Biosciences). Circular consensus sequencing (CCS) was performed on the PacBio Revio platform. Hi-C chromatin conformation capture libraries were prepared from young leaf tissue fixed in 2% formaldehyde under vacuum at ambient temperature for 15 min following a standard Hi-C protocol^82^. Hi-C libraries were sequenced on an Illumina NovaSeq 6000 platform with 150 bp paired-end reads, with a target coverage of approximately 100×.

Total RNA was extracted from fresh leaf tissue using the RNAprep Pure Plus Kit (Tiangen, Beijing, China). mRNA was enriched using poly-T oligo-attached magnetic beads targeting polyadenylated transcripts. RNA quality was checked with a NanoPhotometer (IMPLEN), and RNA concentration and integrity were evaluated using an Agilent 2100 RNA nano 6000 assay kit (Agilent Technologies). RNA libraries were sequenced on the Illumina NovaSeq 6000 platform to generate 150 bp paired-end reads, targeting approximately 6× coverage. Raw RNA-seq reads were processed with Trimmomatic v0.39^83^ for adapter trimming and quality filtering.

### Genome de novo assembly and annotation

To generate chromosome-level assemblies, we first *de novo* assembled contigs from HiFi reads using hifiasm v0.18.6-r513^84^. For samples with Hi-C data, Hi-C reads were mapped to contigs using Juicer v2.0^85^, and Hi-C contact maps were built with the 3D-DNA v201008 pipeline^86^. Chromosome boundaries and assembly misjoins were manually inspected and corrected in Juicebox v1.11.08^87^. For samples lacking Hi-C data, contigs were anchored to pseudochromosomes from conspecific or closely related reference genomes using RagTag v2.0.1^88^. All scaffolded assemblies were manually curated to resolve erroneous inversions, reorder contigs, and remove misassembled contigs where appropriate. Final chromosome sequences were subsequently oriented and renamed to match the chromosome order and orientation of published *Quercus* reference genomes to facilitate consistent cross-species collinearity visualisation. Genome assembly completeness was evaluated using BUSCO v5.4.3^89^ with the embryophyta_odb10 dataset. Consensus quality value (QV) was estimated from HiFi reads using Merqury v1.3^90^. For collinearity analysis across *Quercus* genomes, we aligned all pairs of assemblies using minimap2 v2.21-r1071^91^, identified syntenic blocks and structural variants with SyRI v1.6^92^, and visualised these features using plotsr v1.1.1^93^.

To annotate repetitive regions, tandem repeats were detected using TRF v4.0^94^, and transposable elements (TEs) were annotated with HiTE v3.2^95^. HiTE integrates homology-based and *de novo* approaches to construct a custom repeat library and identify full-length TEs. For protein-coding gene prediction, we conducted *ab initio* gene prediction with BRAKER3 v3.0.8^96^, incorporating GeneMark-ETP and AUGUSTUS v3.3.3^97^, guided by RNA-seq and homologous protein evidence. RNA-seq reads were mapped to each genome using HISAT2 v2.2.1^98^, and transcripts were assembled with StringTie v2.2.1^99^. Training evidence included protein sequences from *Arabidopsis thaliana* Araport11^100^and gene models from published oak genomes. The PASA pipeline v2.5.3^101^ was applied to refine structural annotations, including untranslated regions (UTRs)and alternative splicing isoforms. This pipeline integrated *de novo* and genome-guided transcript assemblies generated by Trinity v2.15.2^102^. For the 19 published assemblies, we predicted protein-coding genes using Helixer v0.3.6^103^, a deep-learning *ab initio* predictor. The number and length of predicted genes are comparable to previous reports. The longest isoform of each predicted gene model was retained to generate a non-redundant gene set, and the completeness of gene models was evaluated using BUSCO v5.4.3^89^.

### Centromere prediction and assessment

For each *Quercus* genome, we searched for *QuCEN146* tandem repeats using Tandem Repeat Annotation and Structural Hierarchy (TRASH) pipeline^104^. A *QuCEN146* monomer were used as the search template. This monomer template was derived from the consensus sequence of *QuCEN146* repeats identified from the *Q. serrata* accession QsXY02, which was generated with Centromics software (https://github.com/ShuaiNIEgithub/Centromics) using HiFi reads. Monomers with length between 141 and 151 bp were classified as canonical *QuCEN146* satellites. To identify candidate centromeric regions, we calculated a weighted CEN-score in 200 kb sliding windows across each chromosome, following the method described in ref^105^. In each window, base pairs overlapping LTR retrotransposons, tandem repeats (*QuCEN146*) and genes were assigned weights of 2.5, 2.5 and -1.0, respectively. These scores were then divided using the Otsu method, which identifies a threshold that separates values into two classes while minimizing intra-class variance. The highest scoring window overlapping a centromeric satellite, or the highest scoring window for species with LTR-rich centromere, was defined as a seed region.

All windows above the threshold flanking this seed region were aggregated to define centromere-proximal intervals. For species with satellite-rich centromeres, candidate centromeric regions were restricted to tandem repeat arrays within the centromere-proximal intervals. *QuCEN146* arrays were defined with a maximum inter-monomer spacing of 100 kb and a minimum copy number of 150 for *QuCEN146*-rich species (> 1 million copies) or 50 copies for *QuCEN146*-poor species (< 1 million copies). Within the centromere-proximal intervals of *QuCEN146*-rich species, adjacent arrays separated by ≤ 1 Mb were merged and treated as candidate centromeric regions. For species with LTR-rich centromeres, precise centromere boundaries were determined through visual inspection of sequence identity dotplots to identify regions of elevated transposon enrichment. We also tested an alternative weighting scheme (LTR =2.5, tandem repeat = 1, gene = -0.5), and obtained qualitatively consistent results with lower spatial resolution (data not shown). As an independent cross-validation, CentIRE v3.0^64^ was applied to the same assemblies to predict centromere intervals based on repeat enrichment, and the concordance between CEN-score-defined and CentIRE-defined boundaries was evaluated.

To assess the assembly continuity of candidate centromeric regions, we calculated GCI^33^ scores and AQI^33^ values, which detect potential assembly gaps and structural errors using low-confidence read segments and shear-based comparison information. To screen for assembly collapses, HiFi reads were aligned to their corresponding chromosomes using pbmm2 v1.9.0 (https://github.com/PacificBiosciences/pbmm2), and alignments were filtered with samtools v1.9^106^. The frequency of the first and second most common bases at each position across the genome was determined using NucFreq v0.1^107^.

### CENH3 ChIP-seq and enrichment analysis

To identify functional centromeres in *Quercus*, we performed CENH3 ChIP–seq for the two representative species: *Q. serrata* (subgenus *Quercus*) and *Q. variabilis* (subgenus *Cerris*). For each species, fresh leaf tissue was collected from the same individuals used for genome assembly (QsLS-3 and JJ; Supplementary Table 1). ChIP was performed as described previously with minor modifications^108^. Briefly, ground fine powder was sufficiently suspended in ice-cold nuclei isolation buffer (M1:10 mM sodium phosphate (pH 7.0), 100 mM NaCl, 0.08% *β*-mercaptoethanol and 12% hexylene glycol) followed by filtration through Miracloth for nuclei preparation. The supernatant was centrifuged at 1,000 × g for 10 min at 4 °C for nuclei collection. The nuclei pellet was washed with nuclei wash buffer (M2: 10 mM sodium phosphate (pH 7.0), 100 mM NaCl, 10 mM MgCl₂, 0.08% *β*-mercaptoethanol, 12% hexylene glycol and 0.5% Triton X-100) for three times. The purified nuclei pellet was resuspended in MNase digestion buffer (MNB: 10% sucrose, 50 mM Tris-HCl (pH 7.5), 4 mM MgCl₂ and 1 mM CaCl₂). Intact chromatin was digested with 1,800 U micrococcal nuclease (MNase) at 37 °C for 10 min followed by adding lysis buffer (1 mM Tris-HCl (pH 7.5), 1 mM PMSF and 1×Complete Mini protease inhibitor). Digested chromatin was assessed by agarose gel electrophoresis. Digested chromatin enriched with mono-, di- and tri-nucleosomal fragments was collected and incubated with 5 μg of the homemade CENH3 antibody at 4 °C overnight. The antibody-chromatin complexes were captured with pre-washed 50 μL rProtein A sepharose beads (GE Healthcare, Cat. 17-1279-03) for 2 h at 4 °C. The incubated beads were sequentially washed with Buffer A containing 50, 100 and 150 mM NaCl for 5 min each. Immunoprecipitated chromatin was eluted using elution buffer (50 mM Tris-HCl (pH 7.5), 10 mM EDTA and 1% SDS) at 65 °C for 15 min. ChIP-enriched DNA was recovered by phenol extraction and cold isopropanol precipitation. Purified ChIP DNA and input DNA were used for library preparation using Rapid Plus DNA Lib Prep Kit V2 (ABclonal, Cat. RK20271). All libraries were sequenced on the Illumina NovaSeq 6000 platform. Two peptides corresponding to the N-terminal region of *Quercus* CENH3 (KHPAHKNPNRKKLSDR and SPRTSPSGRSSQTNRE) were synthesized and used as antigens for antibody production. ChIP-seq experiments performed using both antibodies yielded consistent results; only results generated from the first antibody are presented here.

Paired-end reads from CENH3 ChIP and input libraries were trimmed with fastp v0.23.2^109^ and mapped to the corresponding assembly using Bowtie2 v2.5.4^110^. Read pairs with MAPQ < 10 were discarded using samtools v1.9^106^. PCR duplicates were marked and removed using Picard v3.3.0 (https://github.com/broadinstitute/picard). For reads mapping to multiple loci, the highest-scoring alignment was retained. Alignments with two or more mismatches or unpaired single reads were excluded. Bins per million mapped reads (BPM) coverage values were generated in bedGraph format using bamCoverage tool from deepTools v3.5.6^111^. BedGraph files were converted to 1-based per-base coverage tracks, which were used to compute log_2_(ChIP/Input) enrichment ratios.

### Whole-genome bisulfite sequencing (WGBS) and data analysis

To profile cytosine methylation across *Querucs*, we performed WGBS for 10 *Quercus* species spanning all eight sections of the genus (Supplementary Table 1). All WGBS libraries were prepared from the same individuals used for PacBio HiFi genome sequencing. Genomic DNA was isolated from leaf tissue using the CTAB method. WGBS libraries were constructed following the protocol described in ref^112^ and sequenced on an Illumina NovaSeq 6000 platform, with a target depth of approximately 25×.

Raw WGBS reads were filtered using fastp v0.23.2^109^ and mapped to the corresponding assembly using Bismark v0.24.2^113^. Methylation calls were extracted, and duplicate reads were removed using the standard Bismark workflow. Only cytosine sites supported by at least five reads were retained for methylation quantification. Genome-wide methylation levels were summarised in 10 kb or 200kb non-overlapping windows using CGmapTools v0.1.3^114^. Methylation levels in CG, CHG, and CHH contexts were quantified separately.

### Hi-C chromatin interaction analyses

For the 10 species included in the WGBS analyses, we further characterised genome-wide chromatin interaction frequencies. Hi-C library construction and sequencing are described above. Hi-C sequencing reads were processed using HiC-Pro v3.1.0^115^ to generate normalised chromatin contact maps. Hi-C matrices were converted from HiC-Pro format to GenomicInteractions objects using the hicConvertFormat tool from HiCExplorer v3.7.5^116^. This conversion retained all interaction frequencies and genomic coordinates, restructuring the data for interaction-pair-level queries. Using these converted objects, the genomic distance between each interaction pair was calculated as the absolute difference between the midpoints of the two anchored bins. Interaction pairs were binned into three distance categories: short-range (≤1 Mb), long-range (>10 Mb), and all pairs. For each category, the median contact intensity was computed across all valid interaction pairs per chromosome. These values were used to characterise distance-dependent chromatin decay and to compare interaction patterns between centromeric regions and chromosome arms.

### Annotation and analyses of QuCEN146 tandem repeats

To construct chromosome- and species-level *QuCEN146* consensus sequences, all unique *QuCEN146* monomers from each chromosome or species were aligned using MAFFT v7.526^117^. At each alignment position, the nucleotide present at ≥50% frequency was retained to build the consensus. A global consensus sequence was built by aggregating frequency tables from all chromosomes using the same 50% threshold. Pairwise sequence divergence between chromosome-level consensus sequences was calculated as the proportion of mismatched sites across aligned positions.

To resolve the fine-scale centromeric satellite arrays organisation, we searched for higher-order repeat (HOR) structures within *QuCEN146* arrays. A HOR was defined as an uninterrupted stretch of adjacent monomers where every pairwise comparison had fewer than five mismatches and the block contained at least three monomers. For each centromere, the total HOR count was normalised by the total number of *QuCEN146* monomers in the array to yield the HOR-score^32^. Edit distance between each monomer and its chromosome-specific consensus sequence was calculated using the *stringdist* package in R^118^.

### Centromere haplotype (CenHap) identification

Centromere haplotypes (CenHaps) were identified for each chromosome based on pairwise similarity of *QuCEN146* monomers within centromeric regions. We quantified similarity between homologous chromosomes by counting shared *QuCEN146* monomers (including duplicates). The resulting similarity matrix was clustered with *hcluster* from the *amap* R package and visualised as a heatmap using *pheatmap*. CenHap groups were assigned via visual inspection of these heatmaps. CenHap analyses were restricted to Eurasian white oaks, because deep *QuCEN146* divergence in other *Quercus* lineages prevents robust cross-lineage haplotype comparison. For selected chromosome pairs, sequence identity heatmaps of candidate centromeric regions were generated at 2 kb window resolution using StainedGlass^119^.

### Centromeric LTR retrotransposon characterisation and permutation tests

We classified intact LTR retrotransposons into subfamilies using TEsorter v1.3^120^, with HMM profiles derived from the plant transposable element domain database REXdb-plant^121^. Solo LTRs were identified using the *solo_finder.pl* script from LTR_retriever v2.9.9^122^, with thresholds of ≥ 80% coverage against intact LTRs, alignment score ≥300, and minimum length of 100 bp. LTR insertion times were estimated from sequence divergence between the paired LTRs of individual elements. We applied the formula *T*= (1-LTR identity)/2μ, where μ = 6.75×10^-10^ substitutions per site per year^56^.

To test for spatial associations between centromeric LTRs and *QuCEN146* HOR-scores, we carried out permutation tests across *QuCEN146* arrays from the 46 *Q. serrata* accessions. For each accession, we calculated the mean *QuCEN146* HOR-score in 1-kb windows flanking the start and end coordinates of intact centromeric LTRs, then averaged these values across all accessions. We repeated this procedure for 1,000 sets of randomly placed loci, where random loci were matched to observed LTRs for both count and width distribution per accession.

Observed HOR-scores at LTR positions were compared against the permutation distribution to generate empirical *P*-value. The permutation distribution also defined the significance threshold (α = 0.05) and the null expectation (mean across 1,000 permutations). Permutation tests were performed for the three most abundant centromeric LTR families (*SIRE*, *Retand* and *Reina*) with sufficient sample sizes (*N* > 10). To accommodate chromosome-specific patterns, *Retand* elements were analysed separately for Chr12 versus all other chromosomes (non-Chr12), and *SIRE* elements separately for Chr5 versus all other chromosomes (non-Chr5).

### Phylogenetic clustering of QuCEN146 satellites and LTR retrotransposons

To identify major clades of *QuCEN146* satellite repeats, we analysed non-redundant *QuCEN146* sequences. For each species, we retained unique *QuCEN146* sequences ranked in the top 1% by copy number. These dominant unique sequences collectively constitute ca. 20% of the total *QuCEN146* copies in each satellite-rich species. *QuCEN146* monomers were aligned with MUSCLE v5.3^123^, and approximate maximum-likelihood trees were inferred with FastTreeMP v2.1.11^124^. Local support values for internal branches were calculated using Shimodaira–Hasegawa test.

To resolve phylogenetic relationships between LTRs from centromeric regions and chromosome arms, we performed phylogenetic analyses for three LTR families (*SIRE*, *Retand*, and *Angela*) that predominate in LTR-rich centromeres. This analysis was performed separately for each family within each of the eight species from sections *Cerris* and *Cyclobalanopsis* that possess LTR-rich centromeres. For each family, intact LTR sequences were aligned with MUSCLE v5.3^123^, and approximate maximum-likelihood trees were inferred using FastTreeMP v2.1.11^124^. Branch support for internal nodes was estimated via Shimodaira–Hasegawa tests. We used the GAG domain of *SIRE* elements and the RT domain of *Retand* and *Angela* elements for phylogenetic inference.

### Population genetic analyses

For population genomic analyses of *Q. serrata* and *Q. variabilis*, HiFi reads from each individual were mapped to their respective reference genomes (QsXY02 for *Q. serrata* and SDTS_1 for *Q. variabilis*) using pbmm2 v1.9.0. Genotypes were called with DeepVariant v1.3.0^125^, and individual gVCFs were merged into a multiple-sample VCF using GATK v4.1.8.1^126^. Variants within repetitive regions were excluded. We retained high-quality biallelic SNPs after stringent filtering in VCFtools v0.1.16^127^: sites with mean sequencing depth < 5 or > 100 were removed, sites within repetitive regions were excluded, and all indels were discarded. To investigate population structure within each species, we conducted PCA analyses in PLINK v1.90^128^, based on common SNPs subjected to LD pruning (--indep-pairwise 50 5 0.2), missing rate <10%, and minor allele frequency ≥0.05.

We estimated nucleotide diversity (π) and Tajima’s *D*^129^ within genetic groups of *Q. serrata* and *Q. variabilis*, as well as relative genetic differentiation (*F*_ST_)^130^ and absolute sequence divergence (*d*_XY_)^131^ between groups. All statistics were calculated in 10 kb non-overlapping windows using pixy v1.2.5^132^. For π and *d*_XY_, only windows with ≥1,000 callable sites were retained; for *F*_ST_ and Tajima’s *D*, windows required ≥10 SNPs and ≥ 1,000 callable sites. The population-scaled recombination rate (ρ = 4*N*_e_c) between each pair of SNPs was estimated using LDHELMET v1.9^133^ in a sliding window of 50 SNPs, and estimates were weight-averaged into 10 kb windows. This analysis included SNPs with missing rate < 50% and MAF >0.05, final windows required ≥ 10 SNPs. Identical-by-descent (IBD) blocks were inferred using BEAGLE v4.1^134^.

### Trans-centromeric introgression analyses

To identify signals of trans-centromeric introgression, we compared phylogenetic trees built from centromere-flanking regions (CEN-flank) against those from chromosome arms. For each chromosome, SNPs within 500 kb flanking the centromeric region were used to build a CEN-flank tree, while 10,000 randomly sampled SNPs from the chromosome arms were used for the chromosome-arm tree. SNP matrices were converted to PHYLIP format using vcf2phylip^135^, and maximum likelihood trees were inferred with IQ-TREE2^136^ under the parameters “-T AUTO -m MFP+ASC -B 1000 - bnni -redo”. Only SNPs with missing rate <10% and MAF >0.05 were used for tree inference. Phylogenetic analyses were performed for all *Q. serrata* individuals and other species from section *Quercus*, with *Q. variabilis* as the outgroup.

We validated candidate signals of trans-centromeric introgression for selected cases using efficient local ancestry inference (ELAI) analyses^137^ and the *f*_dm_ statistic^138^. For ELAI, three-way admixture runs were conducted using 15 lower-layer clusters, 20 expectation-maximization (EM) iterations, and 100 admixture generations per optimization. Three replicate runs with distinct random seeds were performed, and ancestry estimates were averaged across replicates. The *f*_dm_ statistic was calculated using the pipeline developed by Simon Martin (https://github.com/simonhmartin/genomics_general). Positive *f*_dm_ values indicates gene flow between P3 and P2, whereas negative value indicates gene flow between P3 and P1. For the illustrative case of introgression between QsKYS03 and *Q. mongolica*, local allele dosages for QsKYS03 were estimated with three ancestry sources: a western *Q. serrata* individual (QsEMS08), *Q. mongolica*, and *Q. variabilis*. The corresponsing *f*_dm_ test used the four-taxon topology (((P1, P2), P3), P4), where P1 = QsKYS03, P2 = QsEMS08, P3 = *Q. mongolica*, and P4 = *Q. variabilis*.

## Data availability

The raw sequencing data generated in this study have been deposited in the Genome Sequence Archive (GSA) under BioProject accessions PRJCA076464 (CRA052185, CRA052709, CRA052805, CRA052817 and CRA053441), PRJCA077317 (CRA052184 and CRA052794), and PRJCA077343 (CRA052177 and CRA052795). The corresponding genome assemblies and gene annotations have been deposited in Zenodo (https://doi.org/10.5281/zenodo.23904349).

## Acknowledgements

We thank Y.-Q. Guo for providing photo of *Q. serrata* cupule. We thank Y. Gao, M. Deng, N. Wang, Z.C Jin for their contribution to sample collection. This work was supported by National Natural Science Foundation of China (32161123003 and NSFC 32301617), Guangdong Flagship Project of Basic and Applied Basic Research (2023B0303050001), Key-Field Research and Development Program of Guangdong Province (2022B1111230001), and Guangdong Science and Technology Plan Project (2023B1212060046).

## Author contributions

Conceptualization: Q.-Q.L., Y.-Y.L., Z.-K.C., W.-L.Z., B.W.

Project administration: A.H., P.S.M., D.E.S., P.S.S., J.C.-B., K.-P.M., Z.-K.C., W.-L.Z., B.W.

Data collection: Y.-H.R., H.-L.Y., B.-F.Z., S.-J.L., Z.S., J.-S., J.-W.N., W.-J.L., L.-J.Q., S.-C.S., L.E., A.T., C.F., Y.S., Y.L., N.-L.Z.,

Analyzed data: Q.-Q.L., Y.-Y.L., Y.-H.R., H.-L.Y., B.-F.Z., B.W.

Writing – original draft: Q.-Q.L., Y.-Y.L., B.W.

Writing – review & editing: R.M., X.-J.L., A.H., P.S.M., J.C.-B., P.K.I., I.H., D.E.S., P.S.S.

All authors read and approved the final version of paper.

